# A meta-analysis of effects of feeding seaweed on beef and dairy cattle performance and methane yield

**DOI:** 10.1101/2021.03.11.434923

**Authors:** Ian J. Lean, Helen M. Golder, Tianna M. D. Grant, Peter J. Moate

## Abstract

There has been considerable interest in the use of red seaweed, and in particular *Asparagopsis taxiformis*, to increase production of cattle and to reduce greenhouse gas emissions. We hypothesized that feeding seaweed or seaweed derived products would increase beef or dairy cattle performance as indicated by average daily gain (ADG), feed efficiency measures, milk production, and milk constituents, and reduce methane emissions. We used meta-analytical methods to evaluate these hypotheses. A comprehensive search of Google Scholar, Pubmed and ISI Web of Science produced 14 experiments from which 23 comparisons of treatment effects could be evaluated. Red seaweed (*A. taxiformis*) and brown seaweed (*Ascophyllum nodosum*) were the dominant seaweeds used. There were no effects of treatment on ADG or dry matter intake (DMI). There was an increase in efficiency for feed to gain by 0.41 ± 0.22 kg per kg [standardized mean difference (SMD) = 0.70 ± 0.35; P = 0.001], but not for gain to feed (P = 0.215), although the direction of the change was for improved efficiency. The type of seaweed used was not a significant covariable for ADG and DMI. Milk production was increased with treatment on weighted mean difference (WMD; 1.35 ± 0.44 kg/d; P <0.001); however, the SMD of 0.45 was not significant (P = 0.111). Extremely limited data suggest the possibility of increased percentages of milk fat (P = 0.040) and milk protein (P = 0.001) on DerSimonian and Laird (D&L) WMD evaluation. The limited data available indicate dietary supplementation with seaweed produced a significant and substantial reduction in methane yield by 5.28 ± 3.5 g/kg DMI (P = 0.003) on D&L WMD evaluation and a D&L SMD of −1.70 (P = 0.001); however, there was marked heterogeneity in the results (*I*^*2*^ > 80%). In one comparison, methane yield was reduced by 97%. We conclude that while there was evidence of potential for benefit from seaweed use to improve production and reduce methane yield more *in vivo* experiments are required to strengthen the evidence of effect and identify sources of heterogeneity in methane response, while practical applications and potential risks are evaluated for seaweed use.

## Introduction

There has been considerable interest in the use of red seaweed, and in particular *Asparagopsis taxiformis* to increase production of cattle and to reduce greenhouse gas emissions [1, 2]. However, several different seaweeds have been fed to cattle and include brown seaweed (*Ascophyllum nodosum*), and *Saragssum wightii*. A commercial product ‘*Tasco*’ has been developed based on *A. nodosum* [3].

To date, there have been several reviews that have provided qualitative overviews of the production responses and the extent of inhibition of methane emissions when seaweed was included in the diets of beef and dairy cows [4, 5]. However, there has been no comprehensive quantitative review of this subject. Given that studies have evaluated the effects of seaweeds on beef cattle production, on dairy cattle production, and on methane emissions, there is clear potential to evaluate the use of seaweeds in cattle production and methane emissions using meta-analytical methods. We hypothesized that feeding seaweed or seaweed derived products would increase beef or dairy cattle performance as indicated by average daily gain (ADG), feed efficiency measures, milk production, and milk constituents, and reduce methane emissions.

## Materials and methods

### Literature search

A comprehensive search of the English language literature used the US National Library of Medicine National Institutes of Health through PubMed (http://www.ncbi.nlm.nih.gov/pubmed), Google Scholar (http://scholar.google.com/), and the ISI Web of Science (http://apps.webofknowledge.com). The search was conducted on 21 January 2021 and searches were based on the following key words with no limits included: seaweed and cattle. We searched the reference lists of papers obtained to identify other studies. One additional paper was identified from a personal communication.

For Google Scholar, 28,400 citation results occurred, and the screening of papers stopped when 50 sequential citations were not relevant. Whereas only 58 and 55 results occurred from Pubmed and ISI Web of Science, respectively. In one case, the authors of an article were contacted to clarify results and to provide additional information.

### Inclusion criteria

Papers were primarily screened on their citation title by 2 reviewers and secondarily screened based on the full text. Experiments were included in the analysis if they met the following inclusion criteria developed by S*cibus* (Camden, NSW, Australia): were full manuscripts from peer-reviewed journals; experiments were *in vivo* and the animals studied were cattle; the experiments evaluated use of seaweed or seaweed derived products for dietary supplementation of cattle; they were randomized; they had a description of the randomization processes employed; they had appropriate analysis of data; they contained sufficient data to determine the effect size for production outcomes (e.g., the number of cattle or pens in each treatment and control group); they had a measure of effect so that the data were amenable to effect size (ES) analysis for continuous data (e.g., standardized mean difference, SMD); and they had a measure of variance (SE or SD) for each effect estimate or treatment and control comparisons. Studies that could not be adequately interpreted, used purposive and non-representative sampling methods or where authors did not respond to clarify their approach, were excluded. Note, one article was included from the pre-print server for Biology, bioRxiv (https://www.biorxiv.org/).

Fig. 1 depicts a PRISMA diagram [6] of the flow of data collection for the meta-analysis. The PRISMA checklist is provided in S1 File. After the initial search and screening 61 different articles (experiments) were identified and papers without a full text (5) were excluded providing 56 papers that were assessed for eligibility. A total of 42 were excluded for the following reasons: the abstract was in English but the full article was in another language (3 experiments), the experiment was *in vitro* (8 experiments), the article was a review or book chapter (7 articles), the experiment had group feeding resulting in pseudo-replication (2 experiments), the experiment was off topic or had irrelevant outcomes (20 experiments), or the experiment lacked measures of variance (2 experiments). A list of articles excluded with the reason is provided in S1 Table. A total of 14 experiments with 23 treatment comparisons were included in the meta-analysis. A list of the experiments and comparisons included in the meta-analysis is provided in Table 1.

**Table 1.**
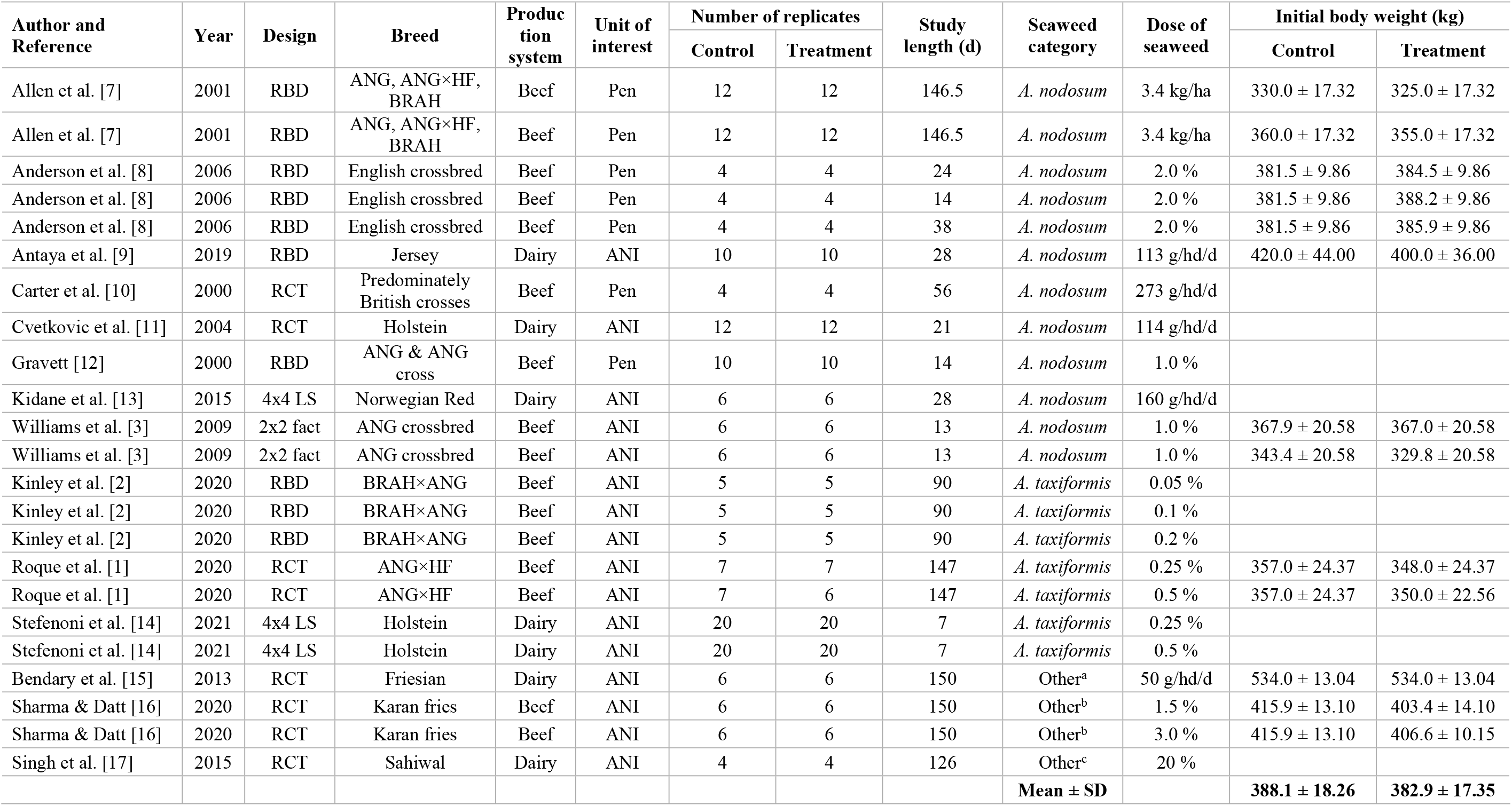
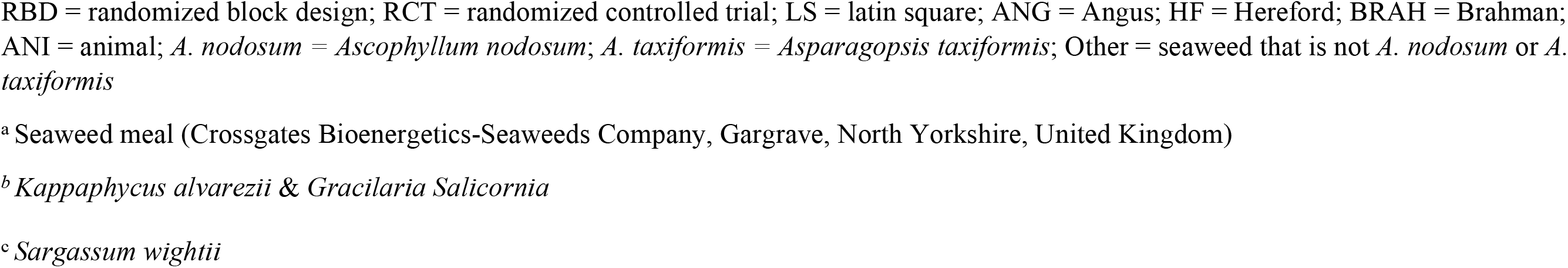
Descriptive information for the comparisons included in the dataset.

| Author and Reference | Year | Design | Breed | Production system | Unit of interest | Number of replicates |  | Study length (d) | Seaweed category | Dose of seaweed | Initial body weight (kg) |  |
| --- | --- | --- | --- | --- | --- | --- | --- | --- | --- | --- | --- | --- |
|  |  |  |  |  |  | Control | Treatment |  |  |  | Control | Treatment |
| Allen et al. [7] | 2001 | RBD | ANG, ANG×HF, BRAH | Beef | Pen | 12 | 12 | 146.5 | *A. nodosum* | 3.4 kg/ha | 330.0 ± 17.32 | 325.0 ± 17.32 |
| Allen et al. [7] | 2001 | RBD | ANG, ANG×HF, BRAH | Beef | Pen | 12 | 12 | 146.5 | *A. nodosum* | 3.4 kg/ha | 360.0 ± 17.32 | 355.0 ± 17.32 |
| Anderson et al. [8] | 2006 | RBD | English crossbred | Beef | Pen | 4 | 4 | 24 | *A. nodosum* | 2.0 % | 381.5 ± 9.86 | 384.5 ± 9.86 |
| Anderson et al. [8] | 2006 | RBD | English crossbred | Beef | Pen | 4 | 4 | 14 | *A. nodosum* | 2.0 % | 381.5 ± 9.86 | 388.2 ± 9.86 |
| Anderson et al. [8] | 2006 | RBD | English crossbred | Beef | Pen | 4 | 4 | 38 | *A. nodosum* | 2.0 % | 381.5 ± 9.86 | 385.9 ± 9.86 |
| Antaya et al. [9] | 2019 | RBD | Jersey | Dairy | ANI | 10 | 10 | 28 | *A. nodosum* | 113 g/hd/d | 420.0 ± 44.00 | 400.0 ± 36.00 |
| Carter et al. [10] | 2000 | RCT | Predominately British crosses | Beef | Pen | 4 | 4 | 56 | *A. nodosum* | 273 g/hd/d |  |  |
| Cvetkovic et al. [11] | 2004 | RCT | Holstein | Dairy | ANI | 12 | 12 | 21 | *A. nodosum* | 114 g/hd/d |  |  |
| Gravett [12] | 2000 | RBD | ANG & ANG cross | Beef | Pen | 10 | 10 | 14 | *A. nodosum* | 1.0 % |  |  |
| Kidane et al. [13] | 2015 | 4x4 LS | Norwegian Red | Dairy | ANI | 6 | 6 | 28 | *A. nodosum* | 160 g/hd/d |  |  |
| Williams et al. [3] | 2009 | 2x2 fact | ANG crossbred | Beef | ANI | 6 | 6 | 13 | *A. nodosum* | 1.0 % | 367.9 ± 20.58 | 367.0 ± 20.58 |
| Williams et al. [3] | 2009 | 2x2 fact | ANG crossbred | Beef | ANI | 6 | 6 | 13 | *A. nodosum* | 1.0 % | 343.4 ± 20.58 | 329.8 ± 20.58 |
| Kinley et al. [2] | 2020 | RBD | BRAH×ANG | Beef | ANI | 5 | 5 | 90 | *A. taxiformis* | 0.05 % |  |  |
| Kinley et al. [2] | 2020 | RBD | BRAH×ANG | Beef | ANI | 5 | 5 | 90 | *A. taxiformis* | 0.1 % |  |  |
| Kinley et al. [2] | 2020 | RBD | BRAH×ANG | Beef | ANI | 5 | 5 | 90 | *A. taxiformis* | 0.2 % |  |  |
| Roque et al. [1] | 2020 | RCT | ANG×HF | Beef | ANI | 7 | 7 | 147 | *A. taxiformis* | 0.25 % | 357.0 ± 24.37 | 348.0 ± 24.37 |
| Roque et al. [1] | 2020 | RCT | ANG×HF | Beef | ANI | 7 | 6 | 147 | *A. taxiformis* | 0.5 % | 357.0 ± 24.37 | 350.0 ± 22.56 |
| Stefenoni et al. [14] | 2021 | 4x4 LS | Holstein | Dairy | ANI | 20 | 20 | 7 | *A. taxiformis* | 0.25 % |  |  |
| Stefenoni et al. [14] | 2021 | 4x4 LS | Holstein | Dairy | ANI | 20 | 20 | 7 | *A. taxiformis* | 0.5 % |  |  |
| Bendary et al. [15] | 2013 | RCT | Friesian | Dairy | ANI | 6 | 6 | 150 | Other<sup>a</sup> | 50 g/hd/d | 534.0 ± 13.04 | 534.0 ± 13.04 |
| Sharma & Datt [16] | 2020 | RCT | Karan fries | Beef | ANI | 6 | 6 | 150 | Other<sup>b</sup> | 1.5 % | 415.9 ± 13.10 | 403.4 ± 14.10 |
| Sharma & Datt [16] | 2020 | RCT | Karan fries | Beef | ANI | 6 | 6 | 150 | Other<sup>b</sup> | 3.0 % | 415.9 ± 13.10 | 406.6 ± 10.15 |
| Singh et al. [17] | 2015 | RCT | Sahiwal | Dairy | ANI | 4 | 4 | 126 | Other<sup>c</sup> | 20 % |  |  |
|  |  |  |  |  |  |  |  |  | **Mean ± SD** |  | **388.1 ± 18.26** | **382.9 ± 17.35** |

RBD = randomized block design; RCT = randomized controlled trial; LS = latin square; ANG = Angus; HF = Hereford; BRAH = Brahman; ANI = animal; *A. nodosum* = *Ascophyllum nodosum*; *A. taxiformis* = *Asparagopsis taxiformis*; Other = seaweed that is not *A. nodosum* or *A. taxiformis*
<sup>a</sup> Seaweed meal (Crossgates Bioenergetics-Seaweeds Company, Gargrave, North Yorkshire, United Kingdom)
<sup>b</sup> *Kappaphycus alvarezii* & *Gracilaria Salicornia*
<sup>c</sup> *Sargassum wightii*

**Fig 1.**
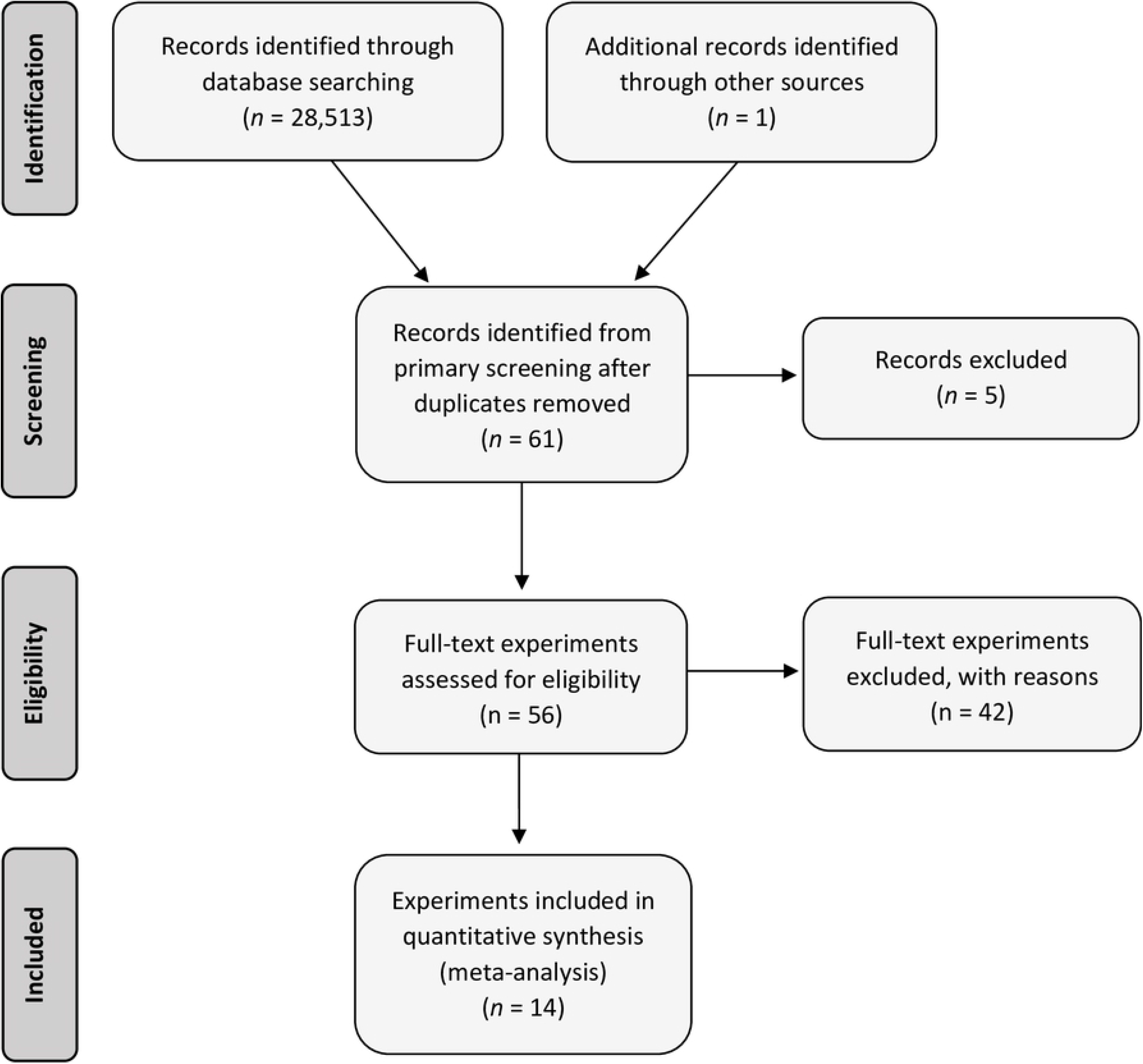
PRISMA flow diagram (adapted from [6]) of the systematic review from initial search and screening to final selection of publications to be included in the meta-analysis on seaweed in cattle.

## Data extraction

All data extracted from each of the experiments that met the inclusion criteria were audited by up to three reviewers. The descriptive data extracted included experiment design, and details about the experiment and the animals used. Design details included the number of: animals or pens, animals/group, and pens/group; experimental and analytical unit (animal or pen). Experimental details included: the number of days on feed, the number of days treatment products were fed, the dose of treatment administered, and diet and delivery methods of product. Animal details included: class of cattle (steers or heifers, or dairy cows), production system (dairy or beef), initial body weight of control and treatment groups, and type of housing and feeding systems. Key descriptive data are provided in Table 1.

Output variables extracted for meta-analysis included: final body weight (FBW, kg), ADG (kg/head/d), dry matter intake (DMI; kg/head/d), gross feed efficiency [ratio of gain to feed (G:F) and ratio of feed to gain (F:G)], milk yield (kg/d), milk fat percentage, milk protein percentage, and methane yield (g/kg DMI) (Tables 2 and 3).

**Table 2.** Mean ± SD of control and treatment group production outcomes for each comparison included in analysis.

| Author and reference | Year | Seaweed category | Final body weight (kg) |  | Average daily gain (kg/d) |  | Dry matter intake (kg/d) |  | Feed to gain (kg/kg) |  | Gain to feed (kg/kg) |  |
| --- | --- | --- | --- | --- | --- | --- | --- | --- | --- | --- | --- | --- |
|  |  |  | Control | Treatment | Control | Treatment | Control | Treatment | Control | Treatment | Control | Treatment |
| Allen et al. [7] | 2001 | *A. nodosum* | 555.0 ± 24.25 | 552.0 ± 24.25 | 1.61 ± 0.10 | 1.63 ± 0.10 | | | 6.70 ± 0.55 | 6.30 ± 0.55 | | |
| Allen et al. [7] | 2001 | *A. nodosum* | 578.0 ± 24.25 | 570.0 ± 24.25 | 1.57 ± 0.10 | 1.55 ± 0.10 | | | 7.20 ± 0.55 | 6.90 ± 0.55 | | |
| Anderson et al. [8] | 2006 | *A. nodosum* | 553.8 ± 17.58 | 552.5 ± 17.58 | 1.45 ± 0.12 | 1.41 ± 0.12 | | | | | 9.14 ± 0.72 | 9.08 ± 0.72 |
| Anderson et al. [8] | 2006 | *A. nodosum* | 553.8 ± 17.58 | 544.4 ± 17.58 | 1.45 ± 0.12 | 1.31 ± 0.12 | | | | | 9.14 ± 0.72 | 9.69 ± 0.72 |
| Anderson et al. [8] | 2006 | *A. nodosum* | 553.8 ± 17.58 | 567.1 ± 17.58 | 1.45 ± 0.12 | 1.52 ± 0.12 | | | | | 9.14 ± 0.72 | 9.02 ± 0.72 |
| Antaya et al. [9] | 2019 | *A. nodosum* | 408.0 ± 35.73 | 392.0 ± 35.73 | | | 18.1 ± 1.26 | 19.3 ± 1.26 | | | | |
| Carter et al. [10] | 2000 | *A. nodosum* | | | 0.86 ± 0.18 | 0.73 ± 0.18 | | | 7.52 ± 3.40 | 5.78 ± 3.40 | | |
| Cvetkovic et al. [11] | 2004 | *A. nodosum* | | | | | 22.7 ± 1.87 | 22.5 ± 1.87 | | | | |
| Gravett [12] | 2000 | *A. nodosum* | | | 1.31 ± 0.16 | 1.36 ± 0.16 | | | 6.23 ± 0.41 | 6.00 ± 0.41 | | |
| Kidane et al. [13] | 2015 | *A. nodosum* | | | | | 18.1 ± 1.15 | 18.1 ± 1.15 | | | | |
| Williams et al. [3] | 2009 | *A. nodosum* | 368.5 ± 19.96 | 365.4 ± 19.96 | 0.05 ± 0.66 | -0.13 ± 0.66 | | | | | | |
| Williams et al. [3] | 2009 | *A. nodosum* | 378.3 ± 19.96 | 364.5 ± 19.96 | 2.68 ± 0.66 | 2.66 ± 0.66 | | | | | | |
| Kinley et al. [2] | 2020 | *A. taxiformis* | | | 1.21 ± 0.38 | 1.24 ± 0.36 | 9.0 ± 1.77 | 8.0 ± 1.14 | 7.45 ± 1.39 | 6.95 ± 0.58 | | |
| Kinley et al. [2] | 2020 | *A. taxiformis* | | | 1.21 ± 0.38 | 1.52 ± 0.29 | 9.0 ± 1.77 | 10.5 ± 1.43 | 7.45 ± 1.30 | 6.60 ± 0.58 | | |
| Kinley et al. [2] | 2020 | *A. taxiformis* | | | 1.21 ± 0.38 | 1.47 ± 0.16 | 9.0 ± 1.77 | 9.4 ± 0.49 | 7.45 ± 0.38 | 6.42 ± 0.58 | | |
| Roque et al. [1] | 2020 | *A. taxiformis* | 589.0 ± 29.37 | 572.0 ± 29.37 | 1.60 ± 0.16 | 1.52 ± 0.16 | 11.3 ± 0.77 | 10.4 ± 0.77 | | | 0.14 ± 0.03 | 0.15 ± 0.03 |
| Roque et al. [1] | 2020 | *A. taxiformis* | 589.0 ± 29.37 | 587.0 ± 27.19 | 1.60 ± 0.16 | 1.56 ± 0.15 | 11.3 ± 0.77 | 9.7 ± 0.71 | | | 0.14 ± 0.03 | 0.16 ± 0.02 |
| Stefenoni et al. [14] | 2021 | *A. taxiformis* | 642.0 ± 77.37 | 645.0 ± 77.37 | | | 25.3 ± 5.81 | 24.5 ± 5.81 | | | | |
| Stefenoni et al. [14] | 2021 | *A. taxiformis* | 642.0 ± 77.37 | 635.0 ± 77.37 | | | 25.3 ± 5.81 | 23.5 ± 5.81 | | | | |
| Bendary et al. [15] | 2013 | Other | | | | | 17.1 ± 0.42 | 17.2 ± 0.42 | | | | |
| Sharma & Datt [16] | 2020 | Other | 426.9 ± 12.02 | 417.2 ± 13.71 | | | 12.2 ± 0.24 | 11.7 ± 0.19 | | | | |
| Sharma & Datt [16] | 2020 | Other | 426.9 ± 12.02 | 418.4 ± 9.76 | | | 12.2 ± 0.24 | 12.0 ± 0.18 | | | | |
| Singh et al. [17] | 2015 | Other | 338.9 ± 18.70 | 334.2 ± 16.50 | | | 9.3 ± 1.40 | 9.6 ± 0.70 | | | | |
|  |  | **Mean ± SD** | **506.9 ± 28.87** | **501.1 ± 28.54** | **1.38 ± 0.26** | **1.38 ± 0.24** | **15.0 ± 1.79** | **14.7 ± 1.57** | **7.14 ± 1.14** | **6.42 ± 0.95** | **5.54 ± 0.44** | **5.62 ± 0.44** |

*A. nodosum* = *Ascophyllum nodosum*; *A. taxiformis* = *Asparagopsis taxiformis*; Other = seaweed that is not *A. nodosum* or *A. taxiformis*.

**Table 3.** Mean ± SD of control and treatment group milk production and methane outcomes for each comparison included in analysis.

| Author | Year | Seaweed category | Milk volume (kg/d) |  | Milk fat (%) |  | Methane (g/kg DMI) |  |
| --- | --- | --- | --- | --- | --- | --- | --- | --- |
|  |  |  | Control | Treatment | Control | Treatment | Control | Treatment |
| Antaya et al. [9] | 2019 | *A. nodosum* | 14.4 ± 1.90 | 15.2 ± 1.90 | 4.4 ± 0.60 | 4.5 ± 0.60 | 22.6 ± 2.78 | 20.6 ± 2.78 |
| Cvetkovic et al. [11] | 2004 | *A. nodosum* | 33.5 ± 2.08 | 35.3 ± 2.08 | 3.9 ± 0.45 | 3.6 ± 0.45 | | |
| Kidane et al. [13] | 2015 | *A. nodosum* | 15.7 ± 1.57 | 16.0 ± 1.57 | 4.3 ± 0.30 | 4.1 ± 0.30 | | |
| Kinley et al. [2] | 2020 | *A. taxiformis* | | | | | 11.0 ± 1.92 | 10.0 ± 3.85 |
| Kinley et al. [2] | 2020 | *A. taxiformis* | | | | | 11.0 ± 1.92 | 6.8 ± 4.02 |
| Kinley et al. [2] | 2020 | *A. taxiformis* | | | | | 11.0 ± 1.92 | 0.3 ± 0.31 |
| Roque et al. [1] | 2020 | *A. taxiformis* | | | | | 17.5 ± 2.65 | 9.5 ± 2.65 |
| Roque et al. [1] | 2020 | *A. taxiformis* | | | | | 17.5 ± 2.65 | 5.0 ± 2.45 |
| Stefenoni et al. [14] | 2021 | *A. taxiformis* | 40.2 ± 8.59 | 40.0 ± 8.59 | 3.6 ± 0.51 | 3.6 ± 0.51 | 13.9 ± 3.00 | 14.4 ± 3.00 |
| Stefenoni et al. [14] | 2021 | *A. taxiformis* | 40.2 ± 8.59 | 37.6 ± 8.59 | 3.6 ± 0.51 | 3.6 ± 0.51 | 13.9 ± 3.00 | 9.8 ± 3.00 |
| Bendary et al. [15] | 2013 | Other | 12.6 ± 0.44 | 14.1 ± 0.44 | 3.2 ± 0.02 | 3.2 ± 0.02 | 11.0 ± 1.92 | 10.0 ± 3.85 |
| Singh et al. [17] | 2015 | Other | 7.3 ± 2.30 | 8.8 ± 1.50 | 5.3 ± 0.16 | 5.4 ± 0.17 | | |
|  |  | **Mean ± SD** | **23.4 ± 3.64** | **23.9 ± 3.52** | **4.0 ± 0.36** | **4.0 ± 0.36** | **14.8 ± 2.48** | **9.5 ± 2.76** |

*A. nodosum* = *Ascophyllum nodosum*; *A. taxiformis* = *Asparagopsis taxiformis*; Other = seaweed that is not *A. nodosum* or *A. taxiformis*; DMI = dry matter intake

### Statistical analysis

Data were structured to allow a classical meta-analytical evaluation of differences in responses of the experimental groups. Many of the experiments in this analysis used multiple treatment comparisons (nesting), and therefore the data had a hierarchical structure. For this reason, meta-regression using multi-level models was used to evaluate the effects of experiment and treatment by taking into account this hierarchical structure [18-20].

Initial data exploration included production of basic statistics using Stata (Version 16, StataCorp. LP, College Station, TX) to examine the data for errors and to estimate the means and measures of dispersion. Normality of the data was examined for continuous variables, by visual and statistical appraisal.

Stata was also used to analyze differences in responses by SMD analysis which is also called ES analysis. These methods have been published in detail in [21] and [22]. The difference between treatment and reference groups means, which is termed ‘treatment’ in the following description, was standardized using the SD of reference and treatment groups. The SMD estimates were pooled using the DerSimonian and Laird random effects models (D&L) [23] and, in the case of methane yield, with the more conservative Knapp-Hartung method (K-H) [24]. Only random effects models were used, as previous work concluded that when there was uncertainty in the evaluative units caused by clustering of observations, the random effects model was appropriate [25].

Robust regressions models (RR) were produced that account for the nested effect of comparisons within experiment [18] and analysed using “*robumeta*” (Stata) as applied by [26]. The RR were developed to account for the two-stage cluster sampling inherent when the ES estimates are derived from a total of *n* = *k*1 + *k*2 + … + *km* estimates from comparisons that were collected by sampling *m* clusters of experiments, that is several comparison estimates are derived from the same experiment [18]. Hence, sampling *kj* ≥ 1 estimates within the *j*^th^ cluster for *j* = 1,…, *m*. Briefly, in this test the mean ES from a series of experiments is described as follows: In this case, the regression model has only an intercept b1 and the weighted mean has the form:

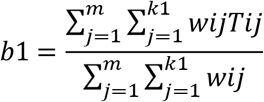

where *m* is the total number of experiments, *k* the total number of comparisons in the extracted database and w*ij* is the weighting for comparisons within experiments and T*ij* is the vector of the ES estimates of comparisons within experiments. If all the estimates in the same experiment are given identical weights, the robust variance estimate (*v*^R^) reduces to:

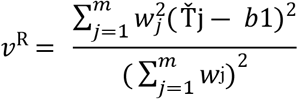

where Ť_j_ is the unweighted mean of the estimates in the j^th^ cluster, b1 is the estimate of the weighted mean, and w_j_ is the total weight given to estimates in the j^th^ cluster. This is a kind of weighted variance which reduces to (m-1)/m^2^ times the variance, when the weights within experiment are identical, and (since the correlation coefficient = 1 in this case) the robust regression standard error equals 1/ m times the variance of Ť_j_ estimated when the weights are equal. Several important aspects of the robust model are highlighted by [18] and the underlying assumptions that; the correlation structure of the T_j_ does not need be known to compute the pooled ES or V^*R*^, only that the vectors of estimates from different experiments are independent and that regularity conditions are satisfied; the experiment or comparison level regressors do not need to be fixed; the theorem is asymptotic based on the number of experiments, rather than the number of comparisons; and the theorem is relatively robust to regularity assumptions.

A random effects weighted mean difference (WMD) between treatment and reference was estimated, with the weighting reflecting the inverse of the variance of the treatments included according to the *nostandard* method in the *metan* model of Stata to allow an interpretation of treatment effects in familiar units (e.g. kg of FBW), rather than ES.

Forest plots were produced for both WMD and SMD results for each outcome variable that incorporated the D&L and RR estimates. The forest plots provided further allow a comparison of *A. taxiformis, A. nodosum*, and ‘*other*’ sources of seaweed evaluated with the D&L and RR methods. Additionally, plots were produced for initial body weight.

Variations among the comparison level SMD were assessed using a chi-squared (Q) test of heterogeneity. Heterogeneity in comparisons reflects underlying differences in clinical diversity of the research site and interventions, differences in experimental design and analytical methods, and statistical variation around responses. The clinical diversity of the site includes all the non-study design aspects of variation, such as facility design, environment, and cattle management that may be measured and controlled for in meta-analysis but are often not reported or measured. Identifying the presence and sources of the heterogeneity improves understanding of the responses to the interventions used. An α level of 0.10 was used because of the relatively poor power of the χ^2^ test to detect heterogeneity among small numbers of trials [27]. Heterogeneity of results among the comparisons was quantified using the *I*^2^ statistic [28]. The *I*^2^ provides an estimate of the proportion of the true variance of effects of the treatment, that is the true variance, tau^2^ (τ^2^) divided by the total variance observed in the comparison [29] that reflect measurement error. Negative values of *I*^2^ are assigned a value of 0, consequently the value *I*^2^ lies between 0 and 100%. An *I*^2^ value between 0 and 40% might not be important, 30 to 60% may represent moderate heterogeneity, 50 to 90% might represent substantial heterogeneity, and 75 to 100% might represent considerable heterogeneity [30]. Both *I*^2^ and τ^2^ are provided to allow readers the opportunity to evaluate both metrics.

A key focus of meta-analysis is to identify and understand the sources of heterogeneity or variation of response among comparisons. However, given the limited number of experiments available the only meta-regression analyses suitable were for category of seaweed intervention for ADG and DMI and production system for DMI.

Presence of publication bias was investigated using funnel plots which are a simple scatter plot of the intervention effect estimates from individual comparisons plotted against comparison precision. The name ‘*funnel plot*’ arises because precision of the intervention effect increases as the size and precision of a comparison increases. Effect estimates from comparisons with a small number of animal units will scatter more widely at the bottom of the graph and the spread narrows for those with higher numbers of units. In the absence of bias, the plot should approximately resemble a symmetrical (inverted) funnel. Funnel plots are available upon request.

## Results and discussion

The literature that was amenable to quantitative review on seaweed use in cattle was reasonably limited with only 14 full texts suitable (Fig 1; Table 1). The experiments used were all published after the year 2000, indicating that they are relatively current. Although these were current some production data indicated only modest production performance (Tables 2 and 3). Funnel plots produced indicated that publication bias was not likely (data not shown). The limited number of comparisons and even fewer experiments limited the type of meta-regressions that could be performed and the use of RR. Only 2 experiments, one on a dairy and one on a beef production system, used Latin Square designs and this precluded evaluation of the effect of study design. As the SD of these were similar to the randomized controlled designs adjustments to the error terms for these were not made.

Differences in FBW were significant for treatment for both RR SMD and RR WMD suggesting that the FBW was lower for treated cattle (Table 4). These findings were not supported by differences in ADG with all models showing little difference in ADG (Table 4; Fig 2). The numerically lower initial body weight for treated cattle supports the contention that FBW differences were substantially influenced by initial BW differences (WMD D&L = −3.08 kg; 95% CI = −7.62 to 1.46; P = 0.183; SMD D&L = −0.28; 95% CI = −0.57 to 0.02; P = −0.57 to 0.02). The comparisons contributing to the observations on FBW and ADG differ but had considerable overlap as 9 comparisons were shared. There was no evidence of difference between *A. taxiformis* and *A. nodosum* interventions on FBW (data not shown) or ADG (Table 4).

**Table 4.** Summary of the meta-analysis using classical meta-analysis methods for the effects of seaweed on production measures. The Table provides the number (N) of experiments and comparisons for each evaluation, the weighted mean difference (WMD) and standardized mean difference (SMD) using both the DerSimonian and Laird (D&L) and robust regression (RR) methods, and the P-value, estimated heterogeneity (*I*^2^) and comparison and experiment variance (τ^2^) of these estimates when available.

| Measure | N comparisons<br>(N experiments) | Effect<br>(95% CI) | P-value | Heterogeneity<br>($I^2$, %) | Variance<br>($\tau^2$) | Meta-regressions<br>(coefficient ± SE; P-value; $\tau^2$) |
| --- | --- | --- | --- | --- | --- | --- |
| **Final body weight** |  |  |  |  |  |  |
| WMD (D&L; kg) | 15<br>(8) | -6.57<br>(-12.23 to -0.90) | 0.023 | 0 | 0 |  |
| WMD (RR; kg) | 15<br>(8) | -5.71<br>(-11.84 to -0.37) | 0.039 |  | 0 |  |
| SMD (D&L) | 15<br>(8) | -0.23<br>(-0.48 to 0.02) | 0.067 | 0 | 0 |  |
| SMD (RR) | 15<br>(8) | -0.27<br>(-0.52 to -0.02) | 0.041 |  | 0 |  |
| **Average daily gain** |  |  |  |  |  |  |
| WMD (D&L; kg/d) | 14<br>(7) | -0.01<br>(-0.05 to 0.03) | 0.730 | 0 | 0 |  |
| WMD (RR; kg/d) | 14<br>(7) | 0.01<br>(-0.09 to 0.07) | 0.711 | | 0 | *A. taxiformis* compared to *A. nodosum* as reference<br>0.05 ± 0.13; P = 0.726; $\tau^2$ = 0 |
| SMD (D&L) | 14<br>(7) | -0.01<br>(-0.31 to 0.29) | 0.947 | 0 | 0 |  |
| SMD (RR) | 14<br>(7) | -0.03<br>(-0.49 to 0.42) | 0.863 | | 0 | *A. taxiformis* compared to *A. nodosum* as reference<br>0.36 ± 0.50; P = 0.538; $\tau^2$ = 0 |
| **Dry matter intake** |  |  |  |  |  |  |
| WMD (D&L; kg/d) | 14<br>(9) | -0.28<br>(-0.63 to 0.07) | 0.119 | 60.95 | 0.35 |  |
| WMD (RR; kg/d) | 14<br>(9) | -0.33<br>(-0.99 to 0.48) | 0.47 | | 0 | Dairy compared to beef as reference<br>0.76 ± 0.38; P = 0.106; $\tau^2$ = 0<br>*A. nodosum* compared to 'Other' as reference<br>0.54 ± 0.51; P = 0.364; $\tau^2$ = 0<br>*A. taxiformis* compared to 'Other' as reference |
| | | | | | | -0.43 ± 0.77; P = 0.622; $\tau^2$ = 0 |
| SMD (D&L) | 14<br>(9) | -0.31<br>(-0.75 to 0.14) | 0.177 | 59.4 | 0.39 |  |
| SMD (RR) | 14<br>(9) | -0.25<br>(-0.91 to 0.41) | 0.393 | | 0 | Dairy compared to beef as reference<br>0.83 ± 0.75; P = 0.324; $\tau^2$ = 0<br>*A. nodosum* compared to 'Other' as reference<br>0.75 ± 0.78; P = 0.389; $\tau^2$ = 0<br>*A. taxiformis* compared to 'Other' as reference<br>0.14 ± 0.85; P = 0.874; $\tau^2$ = 0 |
| **Feed to gain** |  |  |  |  |  |  |
| WMD (D&L) | 7<br>(4) | -0.41<br>(-0.63 to -0.20) | 0.001 | 0.1 | 0 |  |
| SMD (D&L) | 7<br>(4) | -0.70<br>(-1.01 to -0.31) | 0.001 | 0 | 0 |  |
| **Gain to feed** |  |  |  |  |  |  |
| WMD (D&L) | 5<br>(2) | 0.02<br>(-0.01 to 0.04) | 0.133 | 0 | 0 |  |
| SMD (D&L) | 5<br>(2) | 0.35<br>(-0.21 to 0.92) | 0.215 | 0 | 0 |  |
| **Milk yield** |  |  |  |  |  |  |
| WMD (D&L; kg/d) | 7<br>(6) | 1.35<br>(0.91 to 1.78) | <0.001 | 0 | 0 |  |
| SMD (D&L) | 7<br>(6) | 0.45<br>(-0.11 to 1.09) | 0.111 | 65.1 | 0.39 |  |
| **Milk fat** |  |  |  |  |  |  |
| WMD (D&L; %) | 7<br>(6) | 0.06<br>(0.00 to 0.12) | 0.040 | 7.0 | 0 |  |
| SMD (D&L) | 7<br>(6) | 0.12<br>(-0.49 to 0.78) | 0.703 | 66.2 | 0.41 |  |
| **Milk protein** |  |  |  |  |  |  |
| WMD (D&L; %) | 6<br>(5) | 0.06<br>(0.03 to 0.08) | 0.001 | 20.9 | 0 |  |
| SMD (D&L) | 6<br>(5) | 0.59<br>(-0.14 to 1.33) | 0.113 | 73.8 | 0.56 |  |
| **Methane** |  |  |  |  |  |  |
| WMD (D&L; g/kg DMI) | 8<br>(5) | -5.28<br>(-8.78 to -1.78) | 0.003 | 94.2 | 23.6 |  |
| SMD (D&L) | 8 | -1.70 | 0.001 | 84.0 | 1.61 |  |
|  | (5) | (-2.73 to -0.67) |  |  |  |  |
| SMD (K-H)<sup>a</sup> | 8<br>(5) | -1.94<br>(-3.89 to -0.01) | 0.051 | 84.0 | 3.57 |  |

*A. nodosum* = *Ascophyllum nodosum*; *A. taxiformis* = *Asparagopsis taxiformis*; Other = seaweed that is not *A. nodosum* or *A. taxiformis*; DMI = dry matter intake
<sup>a</sup> Knapp-Hartung method [24]

**Fig 2.**
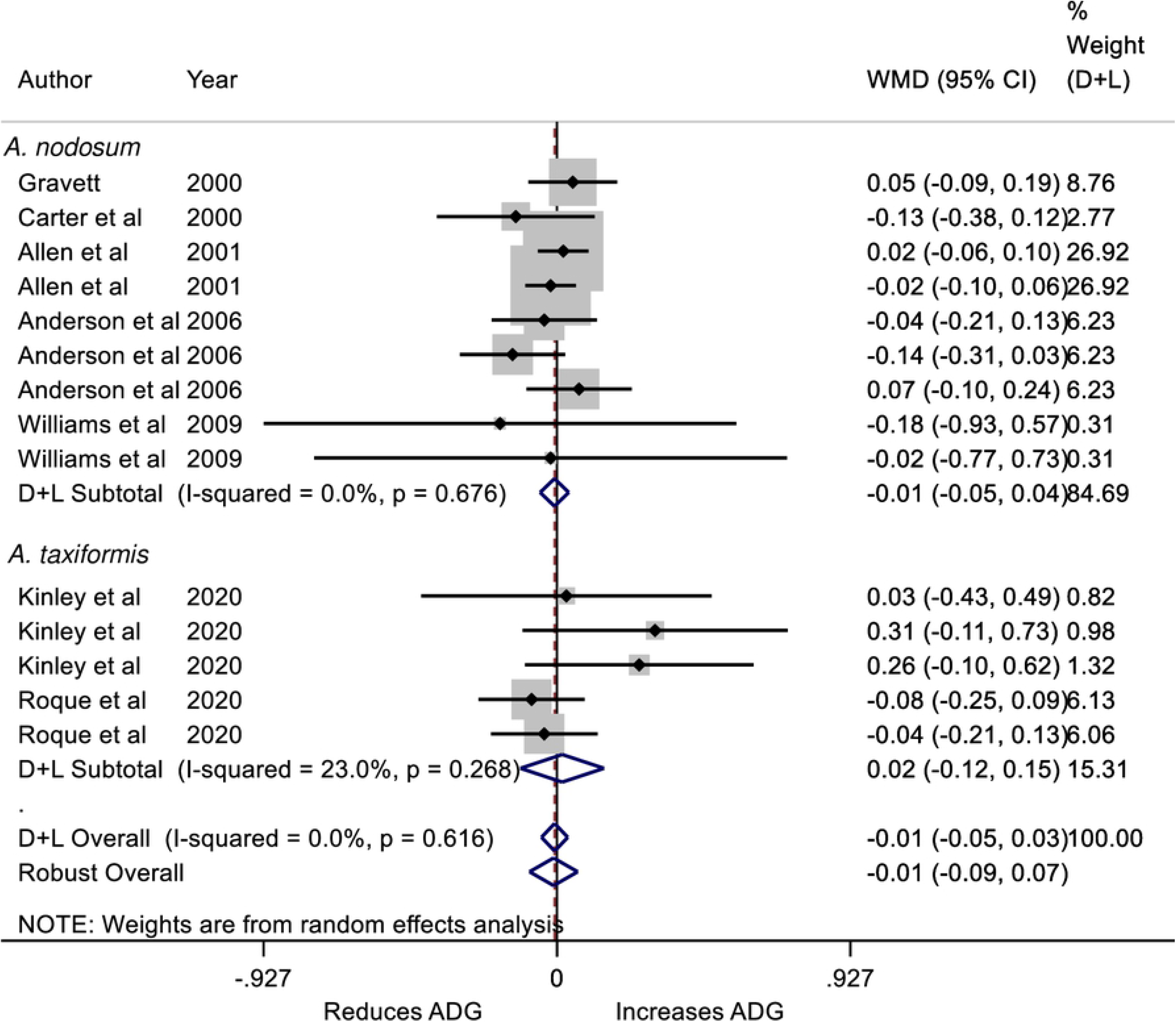
Forest plot of the weighted mean difference (WMD) and 95% CI of the effect of *Ascophyllum nodosum* and *Asparagopsis taxiformis* seaweed intervention on the average daily gain (ADG; kg/d) of cattle. The solid vertical line represents a mean difference of zero or no effect. Points to the left of the line represent a reduction in ADG, while points to the right of the line indicate an increase. Each square around the point effect represents the mean effect size for that comparison and reflects the relative weighting of the comparison to the overall effect size estimate. The larger the box, the greater the comparison contribution to the overall WMD estimate. The weights that each comparison contributed are in the right-hand column. The upper and lower limit of the line connected to the square represents the upper and lower 95% CI for the WMD. The overall pooled WMD and 95% CI pooled using the DerSimonian and Laird (D+L) [23] and robust meta-analytical random effects models [18, 26] are indicated by the respective diamonds at the bottom. The heterogeneity measure, *I*^2^ is a measure of residual variation among comparisons included in the meta-analysis. The ADG was not heterogeneous as indicated by the overall *I*^2^ of 0%.

There was no effect of treatment on DMI (Table 4; Fig 3) and neither the effects of dairy or beef production system nor type of seaweed significantly influenced results (Table 4). Interestingly, these results were heterogenous among comparisons indicting substantial variations in experimental measurement (*I*^*2*^ > 60%; Table 4). The F:G was evaluated in 7- and the G:F in 5-experiments. The F:G was reduced by a significant 0.41 kg per kg with an ES of 0.70 (Table 4); however, these are the less conservative D&L measures as there were insufficient data to evaluate the RR or the effects of differences in seaweed type on F:G. The more limited number of experiments on G:F were not significant (P = 0.215); however, the point direction for the SMD (D&L = 0.35) was consistent with improved feed efficiency from feeding seaweed (Table 4).

**Fig 3.**
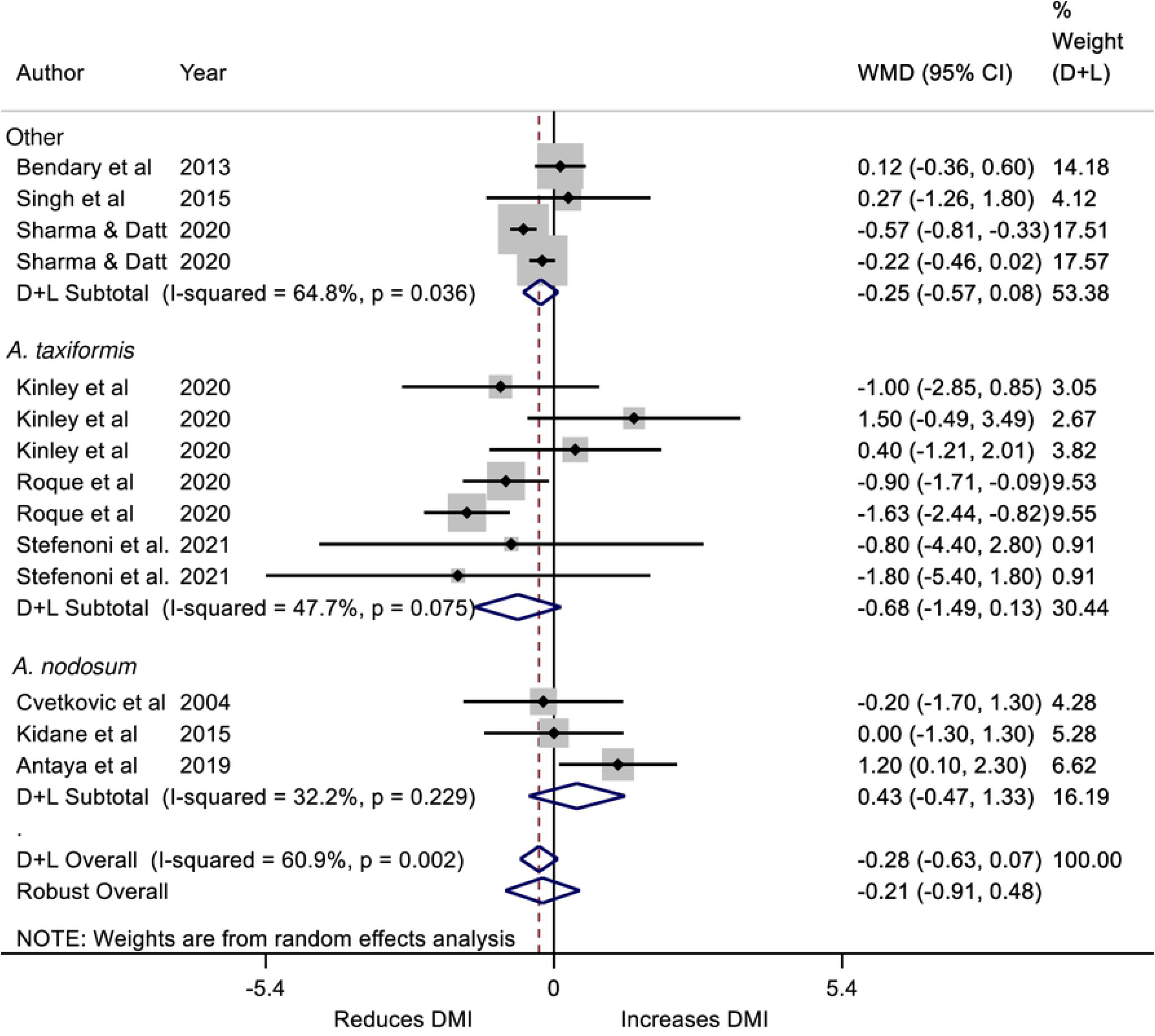
Forest plot of the weighted mean difference (WMD) and 95% CI of the effect of seaweed intervention on the dry matter intake (DMI; kg/d) of cattle. Effects for *Ascophyllum nodosum* and *Asparagopsis taxiformis* and *‘Other’* seaweed interventions are provided as well as an overall effect. The solid vertical line represents a mean difference of zero or no effect. Points to the left of the line represent a reduction in DMI, while points to the right of the line indicate an increase. Each square around the point effect represents the mean effect size for that comparison and reflects the relative weighting of the comparison to the overall WMD estimate. The larger the box, the greater the comparison contribution to the overall estimate. The weights that each comparison contributed are in the right-hand column. The upper and lower limit of the line connected to the square represents the upper and lower 95% CI for the WMD. The overall pooled WMD and 95% CI pooled using the DerSimonian and Laird (D+L) [23] and robust meta-analytical random effects models [18, 26] are indicated by the respective diamonds at the bottom. The heterogeneity measure, *I*^2^ is a measure of residual variation among comparisons included in the meta-analysis. The DMI was substantially heterogeneous as indicated by the overall *I*^2^ of 60.9%.

Milk production was evaluated in only 6 experiments; however, the results were a significant D&L WMD of 1.35 kg/d increase with treatment. However, the D&L SMD of 0.45 was not significant and was heterogenous (*I*^*2*^ = 65.1%; Table 4). There were no significant effects on percentages of milk fat or milk protein on SMD, which were both heterogenous (*I*^*2*^ = 66.2% and 73.8%, respectively). However, the WMD for both milk fat and protein percentages were significantly increased by 0.06% (Table 4). The milk production results contrast with the lack of effect on ADG of treatment, but may be consistent with the efficiency improvement in F:G. The differences in SMD and WMD results reflect sparse data and differences in the weighting between these measures.

There is considerable interest in the potential for *Asparagopsis* to reduce methane emissions and methane yield [1, 2, 14, 31]. The very limited data available for the meta-analysis provide support for the effect to reduce methane yields *in vivo* with a D&L WMD of −5.28 ± 3.5 g/kg of DMI, D&L SMD of −1.70 or K-H SMD of −1.94 indicating a substantial reduction in methane yields. There was marked heterogeneity in the results (*I*^*2*^ > 80%; Table 4; Fig 4). In one comparison the reduction in methane yield with treatment was 97% [2]. These results are consistent with the observations made in *in vitro* studies on the effects of *A. taxiformis* on methane emissions [4] providing further evidence methane emissions is markedly reduced. The mechanism for the reduction in methane emissions and methane yields has been attributed to the bromoform and di-bromochloromethane content of the seaweeds [32, 33] that inhibit methane emissions. However, there are concerns that halogenated gases associated with the bromoforms could cause damage to the ozone layer [4, 34]. At the higher dose of 0.5% inclusion of *A. taxiformis*, [14] found that DMI and milk production and energy corrected milk production were significantly lower than controls and that the milk contained markedly increased concentrations of iodine (> 5 times the control) and bromide (approximately 8 times the control). In the experiment of [14], the concentration of iodine in milk of cows given 0.5% *A. taxiformis* was approximately 3 mg/L, and assuming that a child <3 yr old can drink milk at 1 L/d this is approximately 15 times the upper tolerable intake level [35]. Iodine concentrations in *A. taxiformis* have been reported to range from 8.1 to 11.6 g/kg DM of seaweed [36]. Further, [37] reported that approximately 31% of ingested iodine is transferred to milk indicating there is potential that when cows are fed dietary supplements of *A. taxiformis*, iodine concentrations in milk could be substantially greater than those reported by [14].

**Fig 4.**
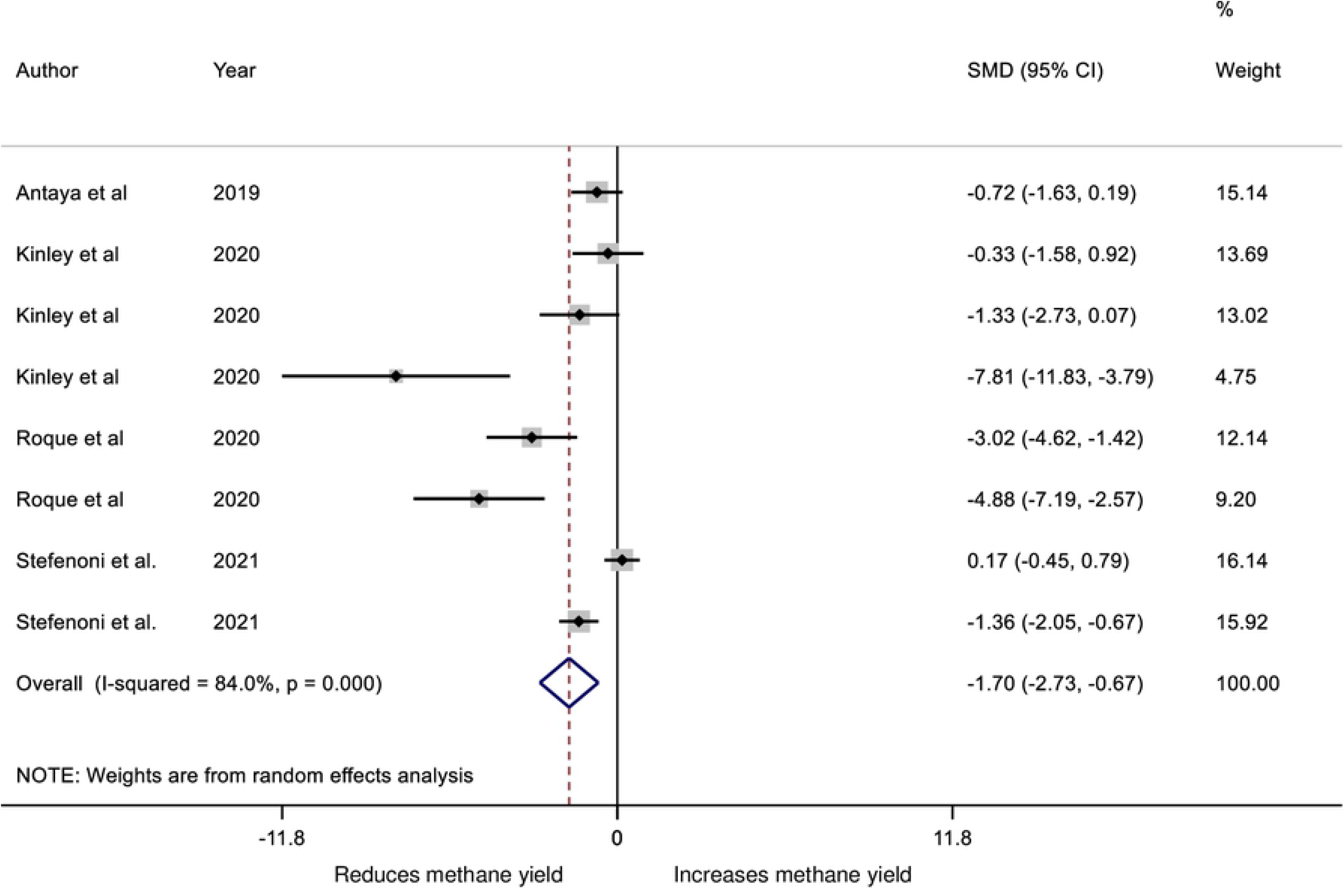
Forest plot of the effect size or standardized mean difference (SMD; standardized using the z-statistic) and 95% CI of the effect of seaweed intervention on methane yield from cattle. The solid vertical line represents a mean difference of zero or no effect. Points to the left of the line represent a reduction in methane yield, while points to the right of the line indicate an increase. Each square around the point effect represents the mean effect size for that comparison and reflects the relative weighting of the comparison to the overall effect size estimate. The larger the box, the greater the comparison contribution to the overall estimate. The weights that each comparison contributed are in the right-hand column. The upper and lower limit of the line connected to the square represents the upper and lower 95% CI for the effect size. The overall pooled effects size or SMD and 95% CI pooled using the DerSimonian and Laird (D+L) [23] and robust meta-analytical random effects models [18, 26] are indicated by the respective diamonds at the bottom. The heterogeneity measure, *I*^2^ is a measure of residual variation among comparisons included in the meta-analysis. Methane yield was considerably heterogeneous as indicated by the overall *I*^2^ of 84.0%.

Although the present analysis indicates that the supplementary feeding of *A. taxiformis* to beef and dairy cattle has some positive effects on animal production and desirable inhibitory effects on methane yields, questions are raised, albeit in a single study, that relate to iodine concentration in *A. taxiformis* and the potential challenges this may bring regarding resultant iodine concentration in milk when feeding *A. taxiformis* to lactating dairy cows.

More *in vivo* experiments are required to strengthen the evidence of production and methane effects in both beef and dairy cows fed under partial mixed ration and pasture-based systems. These studies should use a range of *Asparagopsis* preparations/sources, examine effects on feed intake, and identify sources of heterogeneity in methane response, while practical applications and potential risks are evaluated for seaweed use.

## Acknowledgments

The authors acknowledge Erica Mo for her assistance in data extraction. We thank the authors of [2] for provision of data.

## Supporting information

**SI Table. List of references that were rejected at secondary screening and the reasons**

